# Laboratory evolution can improve algal cell yield and lipid production under mildly cold conditions

**DOI:** 10.1101/2025.08.15.670636

**Authors:** Shu-Yi Lu, Shi-Yu Zhang, Quan-Guo Zhang

## Abstract

One challenge to large-scale microalgae cultivation, e.g., for biodiesel production, is the seasonal low-temperature conditions. We argue that seasonally varying selection in natural environments has prevented algae from better adapting to cold temperatures, and that laboratory evolution offers a promising approach for obtaining cold-adapted algal materials. We conducted a population-level artificial selection experiment with the unicellular green microalgae *Chlorella sorokiniana* at both a benign temperature (25℃) and a mildly cold temperature (15℃). Four artificial selection regimes were established: random selection, selection for high biomass (i.e., cell yield), selection for high lipid production, and rotation between high-biomass and high-lipid selection. We did not observe significant differences among the four selection regimes in evolutionary changes of algal cell yield or lipid yield, suggesting that natural selection at the individual level had dominated the evolutionary changes in our experiment. Compared with the ancestral strain, selection lines that had evolved at 15℃ typically exhibited increased cell yield and reduced lipid content per cell, indicating a trade-off relationship. However, substantial increases in cell yield may compensate for the reduction in lipid content per cell. Notably, three out of 16 selection lines showed > 1-fold increase in cell yield, and one exhibited > 1-fold increase in population-level lipid yield. Selection lines that had evolved at 25℃ displayed even greater increases in both cell and lipid yields, with a positive relationship between cell yield and lipid content per cell. Our results demonstrated the potential for laboratory evolution to obtain algal materials suitable for biofuel production under seasonal low-temperature conditions.

**Highlights:**

- Algae from natural environments not well-adapted to seasonal cold conditions.
- Laboratory evolution under constant conditions with *Chlorella sorokiniana*.
- Both cell yield and lipid production at a low temperature increased.

## 1. Introduction

Microalgae are considered a category of ideal raw materials for biofuel production due to their efficient carbon fixation, rapid growth, strong environmental adaptability, and complete recyclability of biomass (Chen et al., 2018; Chisti, 2007; Patle et al., 2021; Rawat et al., 2013). A large body of research has been devoted to collecting algal materials and identifying culture conditions desirable for faster biomass production and lipid accumulation (Araujo et al., 2011; Guldhe et al., 2019; Sun et al., 2014; Xu & Xiong, 2023; Xu et al., 2024; Zheng et al., 2024). The optimal growth temperature for industrially important microalgae ranges from 20℃ to 30℃ (Ras et al., 2013), hence seasonal low temperatures in non-tropical regions become a hurdle for microalgae cultivation in outdoor environments (Chhandama et al., 2021; Chisti, 2007). Crucially, energy demand is typically higher during low-temperature seasons. Therefore, enhancing algal growth and lipid production under mildly cold conditions (e.g., 10-20℃) would make a significant contribution to clean energy supply.

Collecting algal materials from natural habitats may not be sufficient for obtaining cold-adapted algae. Natural habitats in non-tropical regions typically show seasonal cycles in temperature. Theory suggests that temporally varying environments usually select for niche generalist genotypes with greater geometric mean of fitness across temporal niches (Bell, 2008; Jasmin & Kassen, 2007; Kassen, 2014; Kassen & Bell, 1998). This leads to an expectation that algae from seasonal environments are unlikely to have well adapted to low-temperature conditions. Given the opportunity to evolve in constantly cold conditions, algae may become much better adapted to low temperatures.

Indeed, temporally constant selection pressure may be achieved in the laboratory. The laboratory evolution approach has been used for both answering basic questions in evolutionary biology (Buckling et al., 2009; Elena & Lenski, 2003; Kawecki et al., 2012) and addressing application issues (LaPanse et al., 2021). Examples of the latter included selecting for higher carbon use capacity under elevated CO_2_ conditions (Collins et al., 2006; Collins & Bell, 2004; Jin et al., 2022; Li et al., 2015), and faster carotenoid accumulation under combined blue- and red-light conditions (Fu et al., 2013).

Here we report a laboratory evolution study with *Chlorella sorokiniana*. This unicellular green alga has significant industrial value in biofuel production (Wu et al., 2019). Our study was a population-level artificial selection experiment at two temperatures, 25 ℃ and 15℃, the former a benign temperature and the latter a mildly cold temperature. Artificial selection on the population level has been routinely practiced in conventional crop breeding (Marín, 2021; Murphy et al., 2017). Figure 1A graphically illustrates how to perform directional artificial selection, where random selection is a control treatment. In the present study, we considered two target algal traits, cell yield (a proxy of biomass) and lipid yield. During evolutionary adaptation, improvement in one trait may incur fitness costs in the other (trade-off). Alternatively, positively correlated responses to selection may emerge between two traits (trade-up; Fig. S1). Both trade-off and trade-up relationships are not uncommon (Barrett et al., 2005; Kassen, 2014). We are particularly interested in the possibility of obtaining algal genotypes with both faster biomass production and greater lipid content. As previous studies have suggested that alternating selection targets may actually accelerate evolutionary adaptation (Kashtan et al., 2007) and help to overcome trade-offs between different traits (Zhang et al., 2021), we considered selection regimes with single target traits, as well as a rotational selection scenario (Fig. 1B).

**Fig. 1.**
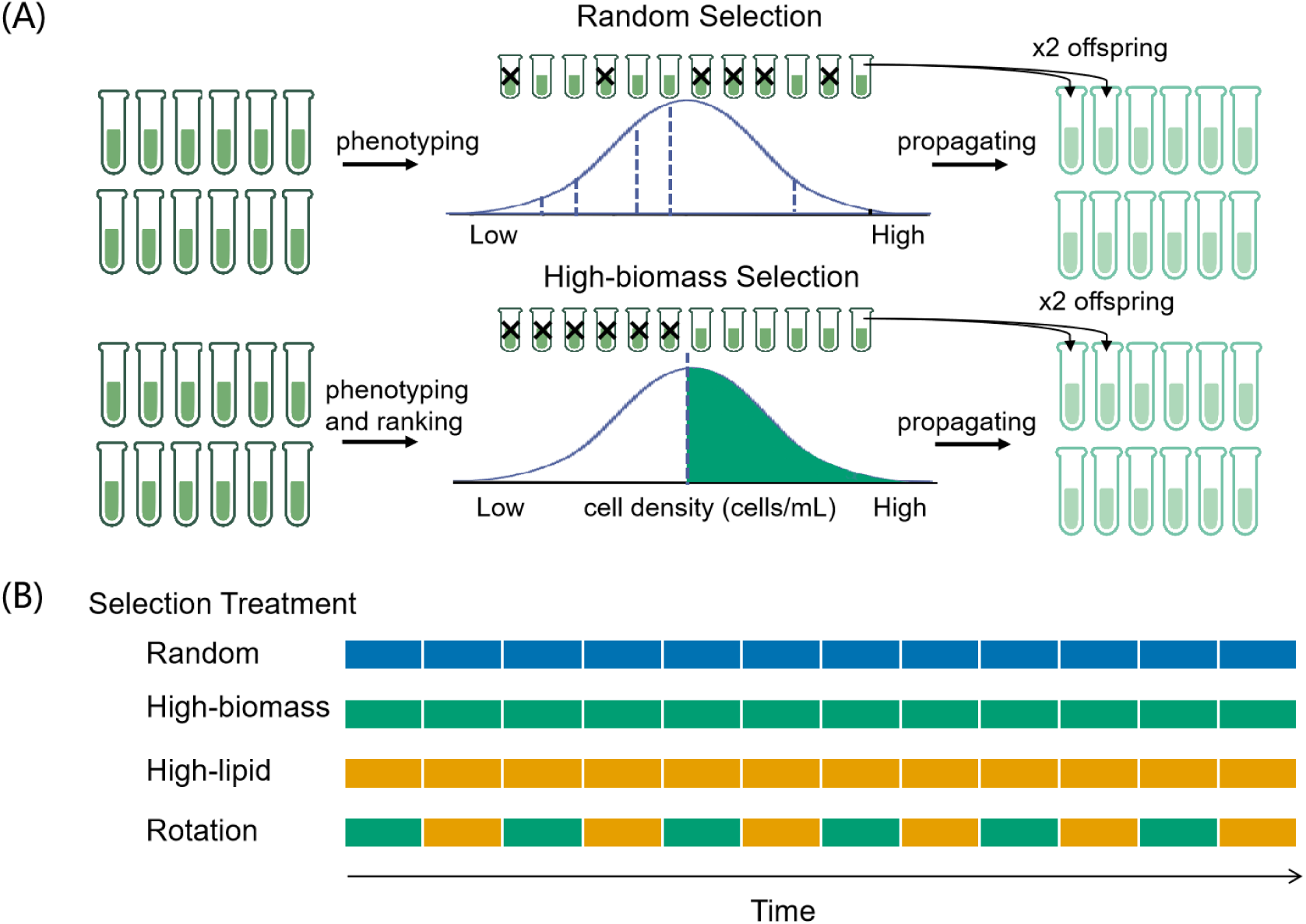
(A) A graphical illustration of the artificial selection protocol. Here one cycle of population propagation and selection was illustrated, with a random selection and a high-biomass selection regimes as examples. (B) Four artificial selection regimes involved in the present study. Constant selection was carried out throughout the experiment under the random, high-biomass, or high-lipid selection regimes, and phases of selection for high-biomass (green) and selection for high lipid (orange) were alternated under the rotational selection regime.

## 2. Materials and Methods

### 2.1 Culture conditions and methods for trait measurements

The *C. sorokiniana* strain FACHB-24 was purchased from the Freshwater Algae Culture Collection of the Chinese Academy of Sciences (FACHB). Algal cultures were grown in BG11 liquid medium under a light: dark cycle of 12 h:12 h, with light intensity of ∼57 μmol photons m^-2^ s^-1^. The batch culture method was used in our experiment. Each culture was propagated in 70 mL of liquid medium in a 100 mL transparent centrifuge tube (with lids loosened to allow for gas exchange), placed in an illumination incubator (GXZ-380A; Ningbo Jiangnan Instrument Factory, China). For every 14 days, 7 mL of each culture was transferred to 63 mL of fresh BG11 medium. The ten-fold population growth during each transfer corresponded to 3.32 generations of binary fission. Cultures were shaken manually once every day, and the positions of the cultures within each incubator layer were alternated daily to minimize environmental heterogeneity among cultures.

Cell density was used as a surrogate of biomass (i.e., cell yield). Specifically, 200 μL of sample from each culture was loaded into 96-well plates, and absorbance at 750 nm was measured using a microplate reader (Bio Tek Synergy H1). Each sample was measured in triplicate. The optical density (OD_750_) was then converted to the cell density. The conversion relationship was as the following: cell density (10^7^ cells mL^-1^) = 4.230 × OD_750_ + 0.039 (Fig. S2). This relationship was established by making a gradient of algal culture dilutions, and measuring their cell densities using a microscope and OD_750_ using a microplate reader.

Lipid content was measured using the sulpho-phospho-vanillin (SPV) colorimetric method (Mishra et al., 2014). Standard lipid stocks were prepared using commercially available soybean oil, and a conversion relationship between lipid content and OD_530_ was established: lipid (mg) = 0.069 × OD_530_ -0.001 (Fig. S3). To measure the lipid content of an algal culture, we centrifuged a 28 mL aliquot for 10 min at 3,176 g (4℃). After the supernatant was discarded, 0.5 mL of concentrated sulfuric acid (98%) was added; then the mixture was heated for 10 min at 90℃, and cooled in an ice bath for 5 min. Subsequently, 3 mL of freshly prepared phospho-vanillin reagent (1.2 g L^-1^, stored in the dark until use) was added. The reaction was carried out in a shaker at 37℃ (200 rpm) for 10 min. Finally, 200 μL of each sample was transferred to a 96-well plate, and OD_530_ was measured. Each sample was measured in triplicate, and the lipid yield (mg L^-1^) was calculated using the lipid standard curve based on algal sample volume.

Under our batch culture conditions, the ancestral algal strain achieved 0.61 ± 0.02 (mean ± S.E.) ×10^7^ cells mL^-1^ of density and 1.51 ± 0.39 mg L^-1^ of lipid yield at 25°C. In the 15°C environment, cell yield was 0.29 ± 0.01 ×10^7^ cells mL^-1^, and lipid yield was 0.55 ± 0.19 mg L^-1^.

### 2.2 The selection experiment

Eight incubators were randomly assigned to the 15℃ or 25℃ temperature conditions, four replicates for each temperature. Within each incubator, one replicate selection line for each of the four artificial selection treatments was set up. Thus, we had a total of 32 selection lines. Algal cultures founded from a single isolate of *C. sorokiniana* was used to start the experiment.

Each selection line consisted of 12 individual algal cultures. The four selection treatments were described as follows. (1) Random selection involved choosing six cultures randomly within a selection line as “parent” populations for the next round of algal propagation, each “parent” culture giving two “offspring” cultures. (2) Under high-biomass selection, the top six cultures in the ranking order of cell yield, measured one day before transferring, were chosen to inoculate the next round of algal cultures. (3) High-lipid selection involved choosing six cultures with greater lipid yield, measured one day before transferring. (4) Under the rotation selection, target traits were switched: high-biomass cultures were chosen at transfer 1, 3, 5, etc., and high-lipid cultures were chosen at transfer 2, 4, 6, etc. (Fig. 1B).

The selection experiment lasted for 12 transfers. During the evolution experiment, we did not observe changes in cell aggregation behavior or cell sizes. At the end of the experiment, we measured the cell and lipid yields of each culture at their respective evolutionary temperatures, and the mean values from every selection line were used in subsequent analysis. Selection response in cell or lipid yield was calculated as a log response ratio (LnRR, logarithm of the ratio of evolved culture relative to the ancestral strain).

### 2.3 Data analysis

Data were analyzed in the R environment (R 4.4.2). One-way ANOVA was used to analyze the difference in selection response in either cell yield or lipid yield among the four selection treatments. Multivariate analysis of variance (MANOVA) was also carried out which considered selection responses in both cell yield and lipid yield as bound response variables. One-sample *t* tests were used for the difference in selection response values and the expected value of zero. Two-sample *t* tests were employed to analyze the differences between the random-selection control and each of the other three selection treatments. Welch’s t tests or Wilcoxon test was used to analyze the differences in measured phenotypic values between the ancestor and each selection treatment according to whether the data satisfies the assumptions of normal distribution and homogeneity of variance.

## 3. Results and discussion

### 3.1 Selection response and artificial selection regime

Our selection lines underwent 12 transfers of laboratory evolution. Evolutionary changes in cell yield or lipid yield, termed “selection response”, were calculated as a log response ratio (LnRR, logarithm of the ratio of evolved culture relative to the ancestral strain). Algae from the four selection treatments at 15°C did not differ in their selection response in cell yield (ANOVA, *F*_3, 12_ = 0.34, *P* = 0.798) or lipid yield (*F*_3, 12_ = 0.28, *P* = 0.839; Fig. 2A; see Tables S1 for comparisons between the random-selection control treatment and each of the other three selection treatments). MANOVA with selection response in cell yield and lipid yield as bound response variables failed to identify a significant difference among selection regimes either (*F*_6, 24_ = 1.01, *P* = 0.443). Similarly, algae from the 25°C environment showed no significant difference among selection regimes (Fig. 2B; Tables S2; ANOVA of selection response in cell yield, *F*_3, 12_ = 0.10, *P* = 0.959; lipid yield, *F*_3, 12_ = 0.57, *P* = 0.647; MANOVA, *F*_6, 24_ = 0.61, *P* = 0.723). The finding that our four population-level selection regimes did not differ in evolutionary changes of algal cell yield or lipid production suggests that natural selection at the individual level has been the predominant force for evolution in our study. Plausibly, there has not been conflicts between population-level traits under artificial selection (cell yield or lipid production) and individual-level fitness (growth traits that determine competitive ability).

**Fig. 2.**
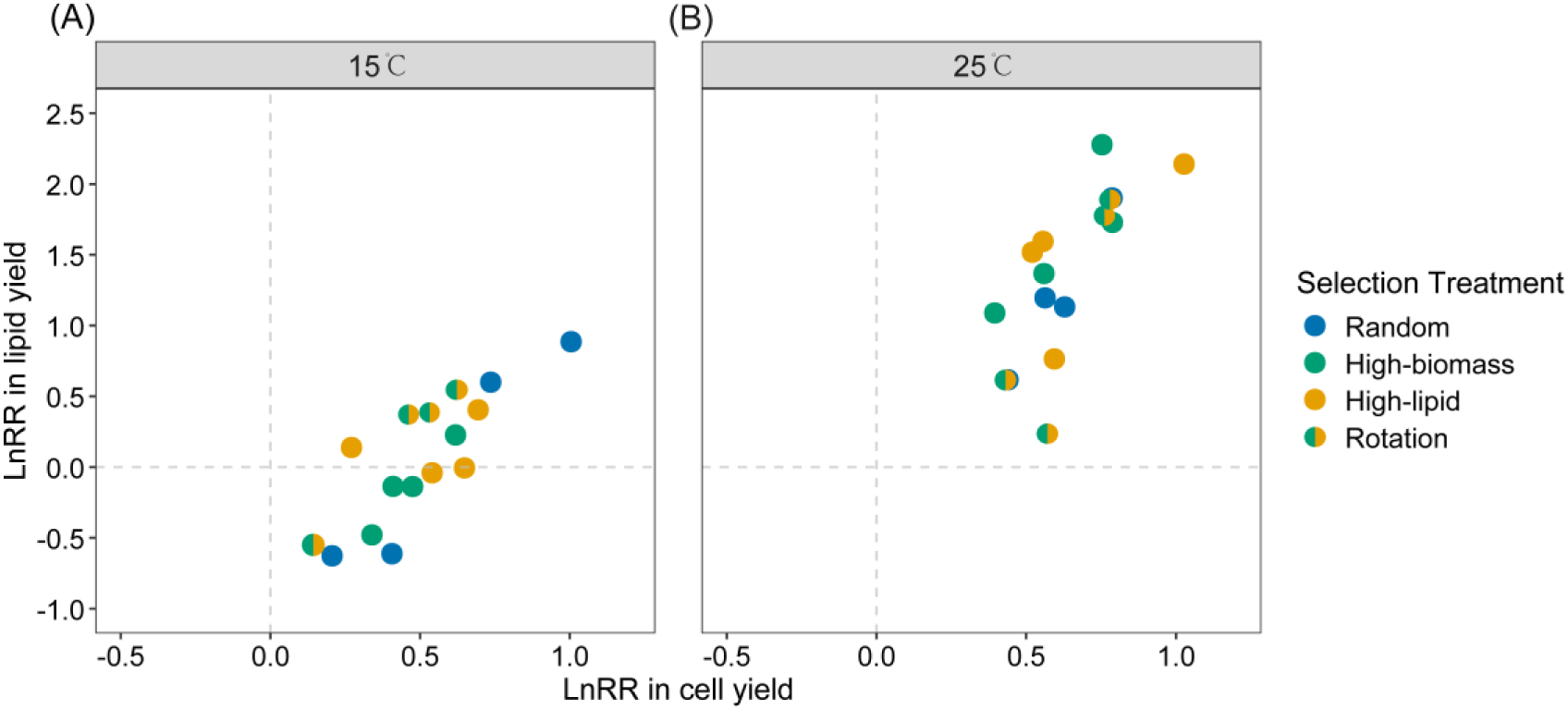
Selection response in cell yield and lipid yield, calculated as the logarithm of the ratio of evolved to ancestral cultures (LnRR). Each scatter represents one single selection line (average values across 12 cultures within each selection line). The horizontal and vertical dashed lines through zero indicated the null hypothesis that evolved cultures did not differ from the ancestral strain.

### 3.2 Selection response and temperature

Out of the 16 selection lines from 15℃, all showed positive values of selection response in cell yield, while eight showed positive selection response values of lipid yield (Fig. 2A). Three selection lines showed > 1-fold of increase in cell yield compared with the ancestral strain (LnRR > 0.693), and one selection line showed > 1-fold of increase in lipid yield. Cultures that had only a small magnitude of evolutionary increase in cell yield typically showed a reduction in lipid yield compared with the ancestral strain (Fig. 2A). The mean value of cell yield selection response was 0.51 ± 0.05, significantly greater than zero (one-sample *t* test, *df* = 15, *P* < 0.001), and mean value of lipid yield selection response was 0.06 ± 0.12, not significantly different from zero (*df* = 15, *P* = 0.317).

All 16 selection lines from 25℃ had positive selection responses in both cell and lipid yields (Fig. 2B), among which six showed > 1-fold increase in both cell and lipid yields compared with the ancestral strain. The mean value of cell yield selection response was 0.63 ± 0.04 (one-sample *t* test, *df* = 15, *P* < 0.001), and the mean value of lipid yield selection response was 1.36 ± 0.15 (*df* = 15, *P* < 0.001).

Algal cultures under both two temperature conditions showed increased cell yield and lipid yield, however, with non-trivial differences. The magnitude of increases in cell and lipid yield was greater under the 25℃ environment (Fig. 2). This may be simply because organisms show faster evolutionary adaptation in warmer conditions, as suggested by the evolutionary speed hypothesis (Bennett & Lenski, 1997; Chu et al., 2020; Fischer, 1960; Rohde, 1992; Sørensen et al., 2018).

Lipid content per cell was also calculated based on cell yield and population-level lipid yield. Selection lines from 15℃ showed a reduction in lipid content per cell compared with the ancestral strain, with a mean value of -0.45 ± 0.08 (*df* = 15, *P* = 1.000; Fig. 3A). A positive selection response in cell yield was accompanied by a reduction in lipid content per cell, which is corresponding to the trade-off scenario as shown in Fig. S1C. This trade-off likely reflects a dilemma in resource re-allocation under cold conditions. Microalgae encounter multiple physiological challenges at low temperatures, including reduced chemical reaction rates, limited enzyme activity, decreased cell membrane fluidity, protein denaturation, and increased water viscosity (Hassan et al., 2016, 2020). Survival under such conditions may be enhanced by the accumulation of major carbon storage compounds such as starch (carbohydrates) and lipids. Starch and lipid metabolism are highly interconnected, sharing glyceraldehyde-3-phosphate (G3P) as a common precursor (Ran et al., 2019; Takeshita et al., 2014). Moreover, the tricarboxylic acid (TCA) cycle and the *de novo* fatty acid synthesis pathway in algae function are competing carbon storage pathways, both utilizing phosphoenolpyruvate (PEP) as a common intermediate metabolite (Zhu et al., 2022). This competition directly affects lipid synthesis and transport. Therefore, cells may prioritize rapid proliferation over energy storage at lower temperatures. Where growth rate is limited by factors other than temperature (e.g., lack of nitrogen, phosphorus, and other nutrient elements), algal cells may instead accumulate lipids at the expense of biomass productivity (Ho et al., 2012; Mandal & Mallick, 2011; Zhu et al., 2022). It is noteworthy that a very large magnitude of increase in cell yield can compensate for reduced lipid content per cell, enhancing population-level lipid production (mg lipid content per L culture; Fig. 2A).

**Fig. 3.**
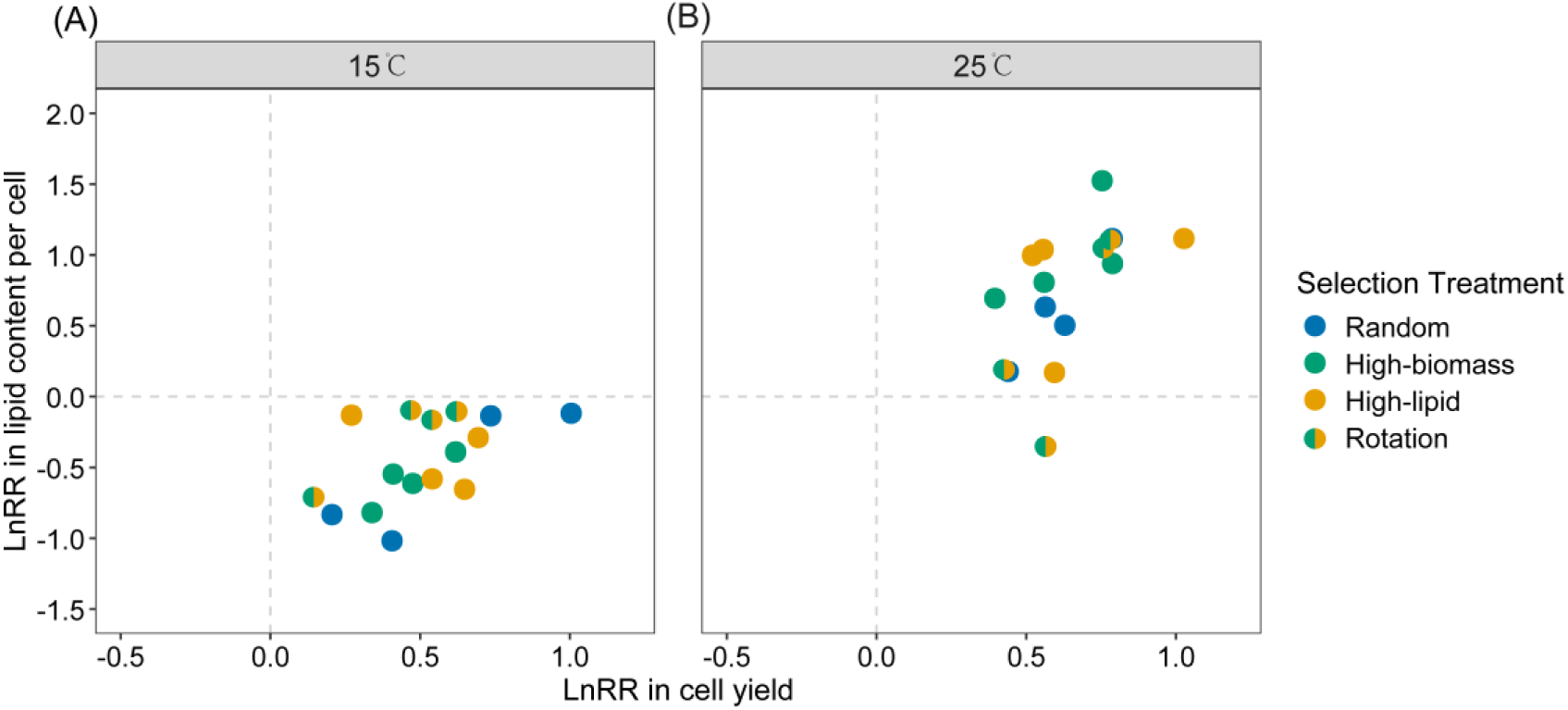
Selection response in cell yield and lipid content per cell. Symbols as in Figure 2.

By contrast, In the 25℃ environment, algal cultures typically showed significant positive selection response in lipid content per cell, with a mean value of 0.73 ± 0.12 (*df* = 15, *P* < 0.001; Fig. 3B). An evolutionary trade-up relationship was observed between cell yield and lipid content per cell in the 25℃ environment, matching the trade-up scenario in Fig. S1A. This suggests that the potential conflict in resource re-allocation discussed above may be mitigated under benign temperature conditions. These results showed that temperature alter the trade-off relationships among organismal traits.

It has been suggested that natural selection in temporally varying environments usually favors niche generalists (Kumawat et al., 2025; Rainey & Travisano, 1998; Wittmann et al., 2023). We expected that laboratory evolution of algal cultures under constant temperature conditions would lead to an improvement in growth performance in those specific conditions. The present study showed that laboratory evolution can increase algal performance in both biomass and lipid production, and evolutionary improvement was even greater in the benign environment (Fig. 2). On average, at both 15°C and 25°C, the cell yield, lipid yield, and lipid content per cell in all evolutionary lines were significantly higher than those of their ancestral strains (Table S3 & S4). Experimental evolution is a promising approach for increasing organism’s tolerance to stresses such as extreme temperature, salinity, and light conditions (Barten et al., 2022), and for enhancing the production of certain metabolites (Cho et al., 2024; Fu et al., 2013; Yu et al., 2013). Here we emphasize that seasonal cold conditions may restrict the industrial applications of microalgae in non-tropical regions, and that laboratory evolution can help to obtain algal materials desirable for biofuel production under mildly cold conditions.

## 4 Conclusions

In this study, laboratory evolution proved to be an effective approach for enhancing algal cell yield and lipid production under mildly cold conditions, although the improvements were more pronounced in warm environments. Temperature was found to influence the trade-offs between biological traits. No significant differences were observed among the four selection regimes in the evolutionary changes of algal cell yield or lipid yield, suggesting that natural selection at the individual level was the predominant driver of these changes. Our results suggested that it pays off for industrial applications of microalgae to use materials specifically adapted to different temperature conditions in different seasons.

## Data availability statement

Data associated with this study are available at figshare (https://doi.org/10.6084/m9.figshare.29319644).

## Acknowledgments

This work was supported by the National Natural Science Foundation of China (32371687).

## Author Contributions

SYL and QGZ designed the study; SYL and SYZ performed experiments; SYL and QGZ analyzed data; all authors wrote the paper.

## Supplementary information

**Figure S1.**
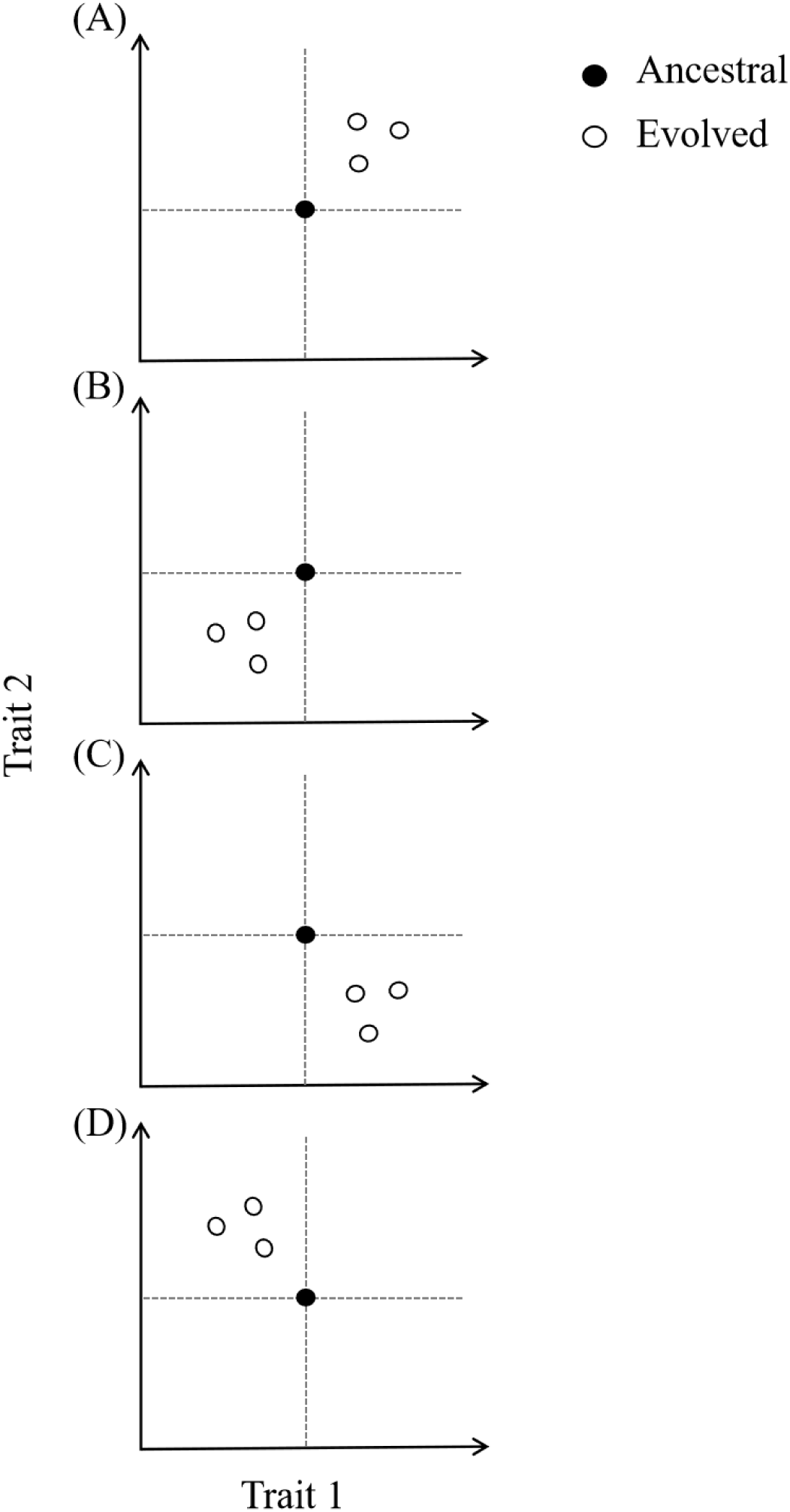
A graphical illustration of evolutionary trade-up (A, B) and trade-off (C, D) relationships between two traits.

**Figure S2.**
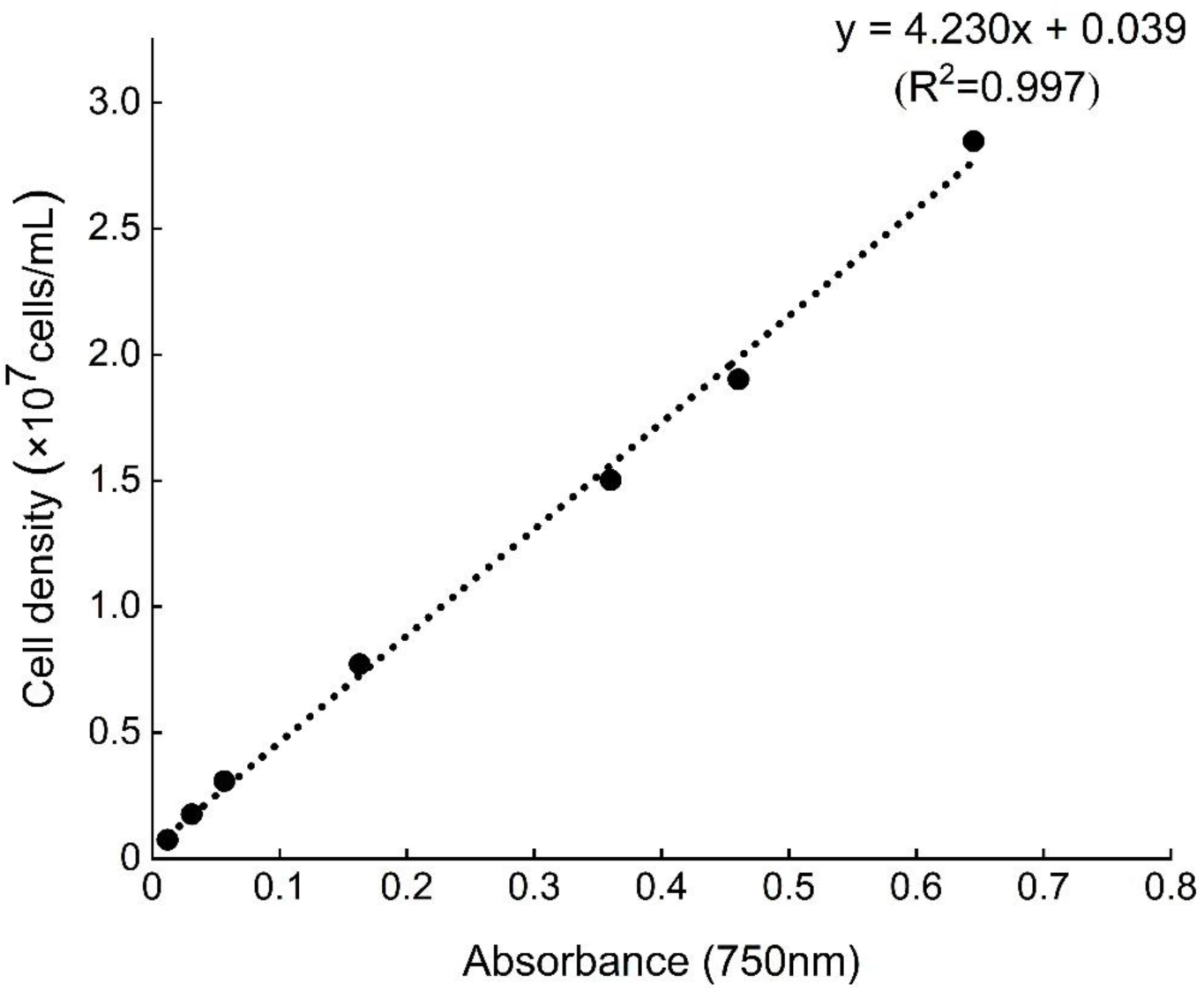
The relationship between optical density (OD_750_) and cell density of *Chlorella sorokiniana*. This relationship was established by making a gradient of algal culture dilutions, and measuring their cell densities using a microscope and OD_750_ using a microplate reader.

**Figure S3.**
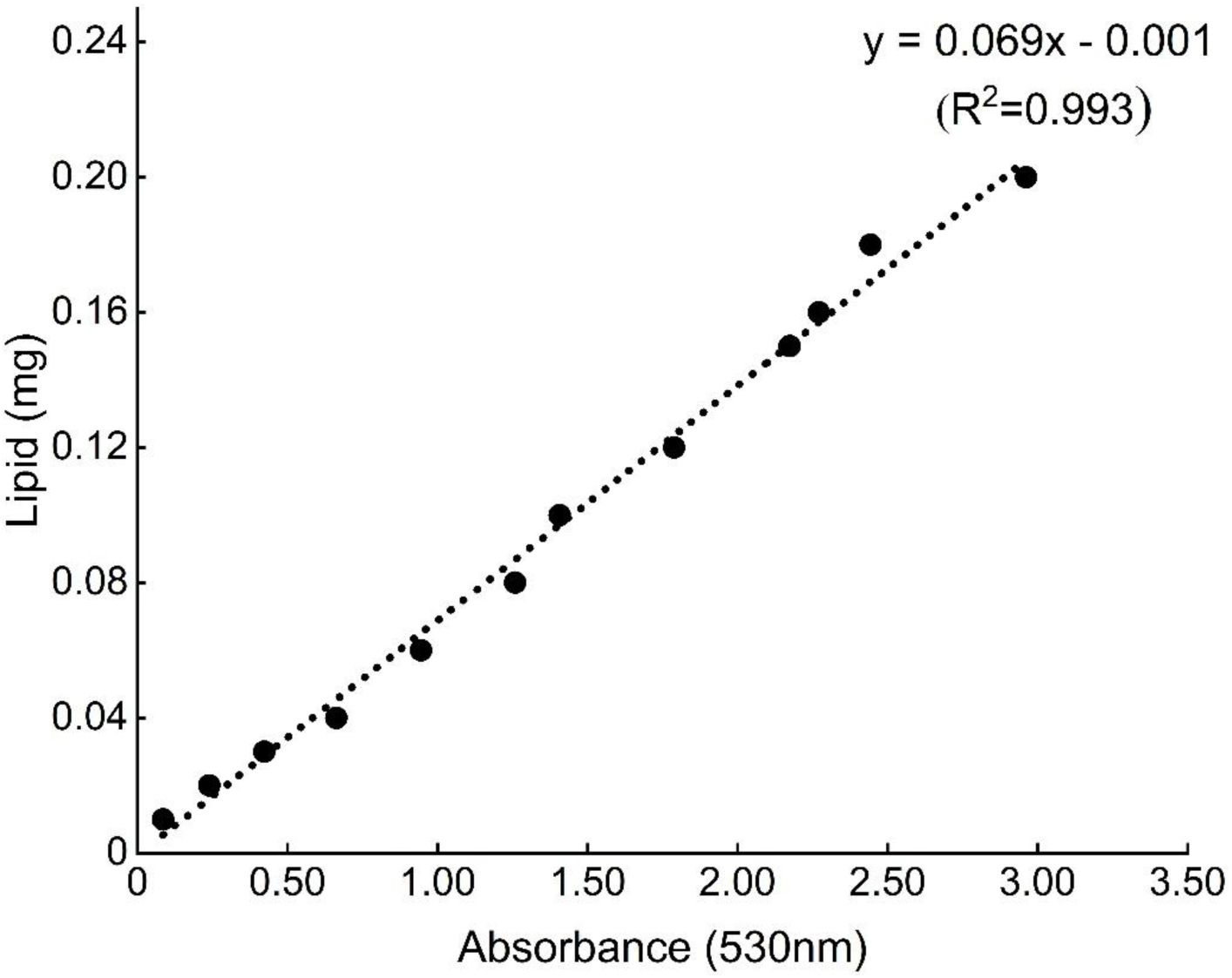
Lipid standard curve was established based on the sulpho-phospho-vanillin (SPV) colorimetric method. Standard lipid stocks were prepared using commercially available soybean oil.

**Table S1.** Summary of the selection response values in the 15°C selection lines. Here are shown the mean (± se) values of selection lines under each selection treatment (n = 4). Differences between the random-selection control and each of the other three treatments were analyzed using two-sample *t* tests.

|  | Cell yield |  | Lipid yield |  | Lipid content per cell |  |
| --- | --- | --- | --- | --- | --- | --- |
| | Mean ± se | Difference from random-selection | Mean ± se | Difference from random-selection | Mean ± se | Difference from random-selection |
| Random | 0.59 ± 0.18 | | 0.06 ± 0.40 | | -0.53 ± 0.23 | |
| High-biomass | 0.46 ± 0.06 | $df = 3.67, P = 0.535$ | -0.13 ± 0.14 | $df = 3.78, P = 0.673$ | -0.59 ± 0.09 | $df = 3.85, P = 0.806$ |
| High-lipid | 0.54 ± 0.09 | $df = 4.60, P = 0.815$ | 0.12 ± 0.10 | $df = 3.39, P = 0.887$ | -0.41 ± 0.12 | $df = 4.54, P = 0.689$ |
| Rotation | 0.44 ± 0.10 | $df = 4.86, P = 0.511$ | 0.17 ± 0.26 | $df = 5.12, P = 0.824$ | -0.27 ± 0.15 | $df = 5.21, P = 0.402$ |

**Table S2.** Summary of the selection response values in the 25°C selection lines. Here are shown the mean (± se) values of selection lines under each selection treatment (n = 4). Differences between the random-selection control and each of the other three treatments were analyzed using two-sample *t* tests.

|  | Cell yield |  | Lipid yield |  | Lipid content per cell |  |
| --- | --- | --- | --- | --- | --- | --- |
| | Mean ± se | Difference from random-selection | Mean ± se | Difference from random-selection | Mean ± se | Difference from random-selection |
| Random | 0.60 ± 0.07 | | 1.21 ± 0.26 | | 0.61 ± 0.19 | |
| High-biomass | 0.62 ± 0.09 | $df = 5.70, P = 0.874$ | 1.62 ± 0.26 | $df = 5.99, P = 0.315$ | 0.99 ± 0.18 | $df = 5.98, P = 0.202$ |
| High-lipid | 0.67 ± 0.12 | $df = 4.97, P = 0.636$ | 1.50 ± 0.28 | $df = 5.97, P = 0.478$ | 0.83 ± 0.22 | $df = 5.90, P = 0.479$ |
| Rotation | 0.63 ± 0.08 | $df = 5.86, P = 0.809$ | 1.12 ± 0.42 | $df = 5.05, P = 0.857$ | 0.48 ± 0.35 | $df = 4.68, P = 0.773$ |

**Table S3.** Summary of the measured values in the 15°C selection lines. Here are shown the mean (± se) values of selection lines under each selection treatment (n = 4). Differences between the ancestor and each selection treatment were analyzed using Welch’s t tests or Wilcoxon test according to whether the data satisfies the assumptions of normal distribution and homogeneity of variance. Welch’s t tests are performed because the response variable cell yield follows a normal distribution but does not meet the assumption of homogeneity of variance. Wilcoxon tests are performed because the response variables lipid yield and lipid content per cell do not meet the assumption conditions.

| | Cell yield ($\times 10^7$ cells mL<sup>-1</sup>) | | Lipid yield (mg L<sup>-1</sup>) | | Lipid content per cell (pg cell<sup>-1</sup>) | |
| --- | --- | --- | --- | --- | --- | --- |
| | Mean ± se | Difference from ancestor | Mean ± se | Difference from ancestor | Mean ± se | Difference from ancestor |
| Ancestor | 0.29 ± 0.01 | | 0.55 ± 0.19 | | 1.86 ± 0.65 | |
| Random | 0.55 ± 0.10 | $df = 3.02, P = 0.076$ | 0.73 ± 0.26 | $W = 11, P = 0.045$ | 1.20 ± 0.26 | $W = 14, P = 0.088$ |
| High-biomass | 0.47 ± 0.03 | $df = 3.19, P = 0.008$ | 0.50 ± 0.07 | $W = 12, P = 0.057$ | 1.05 ± 0.09 | $W = 16, P = 0.130$ |
| High-lipid | 0.51 ± 0.04 | $df = 3.08, P = 0.017$ | 0.63 ± 0.07 | $W = 11, P = 0.045$ | 1.27 ± 0.16 | $W = 15, P = 0.107$ |
| Rotation | 0.46 ± 0.04 | $df = 3.08, P = 0.030$ | 0.71 ± 0.14 | $W = 10, P = 0.035$ | 1.48 ± 0.19 | $W = 13, P = 0.071$ |
| Mean | 0.50 ± 0.03 | $df = 15.94, P < 0.001$ | 0.64 ± 0.07 | $W = 44, P = 0.002$ | 1.25 ± 0.09 | $W = 58, P = 0.008$ |

**Table S4.** Summary of the measured values in the 25°C selection lines. Here are shown the mean (± se) values of selection lines under each selection treatment (n = 4). Differences between the ancestor and each selection treatment were analyzed using Welch’s *t* tests or Wilcoxon test according to whether the data satisfies the assumptions of normal distribution and homogeneity of variance. Welch’s *t* tests are performed because the response variable cell yield follows a normal distribution but does not meet the assumption of homogeneity of variance. Wilcoxon tests are performed because the response variables lipid yield and lipid content per cell do not meet the assumption conditions.

| | Cell yield ($\times 10^7$ cells mL<sup>-1</sup>) | | Lipid yield (mg L<sup>-1</sup>) | | Lipid content per cell (pg cell<sup>-1</sup>) | |
| --- | --- | --- | --- | --- | --- | --- |
| | Mean ± se | Difference from ancestor | Mean ± se | Difference from ancestor | Mean ± se | Difference from ancestor |
| Ancestor | 0.61 ± 0.02 | | 1.51 ± 0.39 | | 2.37 ± 0.55 | |
| Random | 1.13 ± 0.08 | $df = 3.28, P = 0.007$ | 5.64 ± 1.56 | $W = 1, P = 0.003$ | 4.79 ± 0.97 | $W = 4, P = 0.007$ |
| High-biomass | 1.16 ± 0.10 | $df = 3.18, P = 0.012$ | 8.40 ± 2.26 | $W = 0, P = 0.002$ | 7.02 ± 1.46 | $W = 2, P = 0.004$ |
| High-lipid | 1.23 ± 0.16 | $df = 3.07, P = 0.030$ | 7.60 ± 1.98 | $W = 1, P = 0.003$ | 6.02 ± 1.05 | $W = 3, P = 0.005$ |
| Rotation | 1.16 ± 0.10 | $df = 3.21, P = 0.009$ | 5.85 ± 2.06 | $W = 4, P = 0.007$ | 4.72 ± 1.40 | $W = 7, P = 0.017$ |
| Mean | 1.17 ± 0.05 | $df = 18.43, P < 0.001$ | 6.87 ± 0.94 | $W = 6, P < 0.001$ | 5.64 ± 0.61 | $W = 16, P < 0.001$ |

