## Supplementary material for "Laboratory evolution can improve algal cell yield and lipid production under mildly cold conditions": Figure S1, Figure S2, Figure S3, Table S1, Table S2, Table S3, Table S4

**Supplementary information**

**Figure S1** A graphical illustration of evolutionary trade-up (A, B) and trade-off (C, D) relationships between two traits.

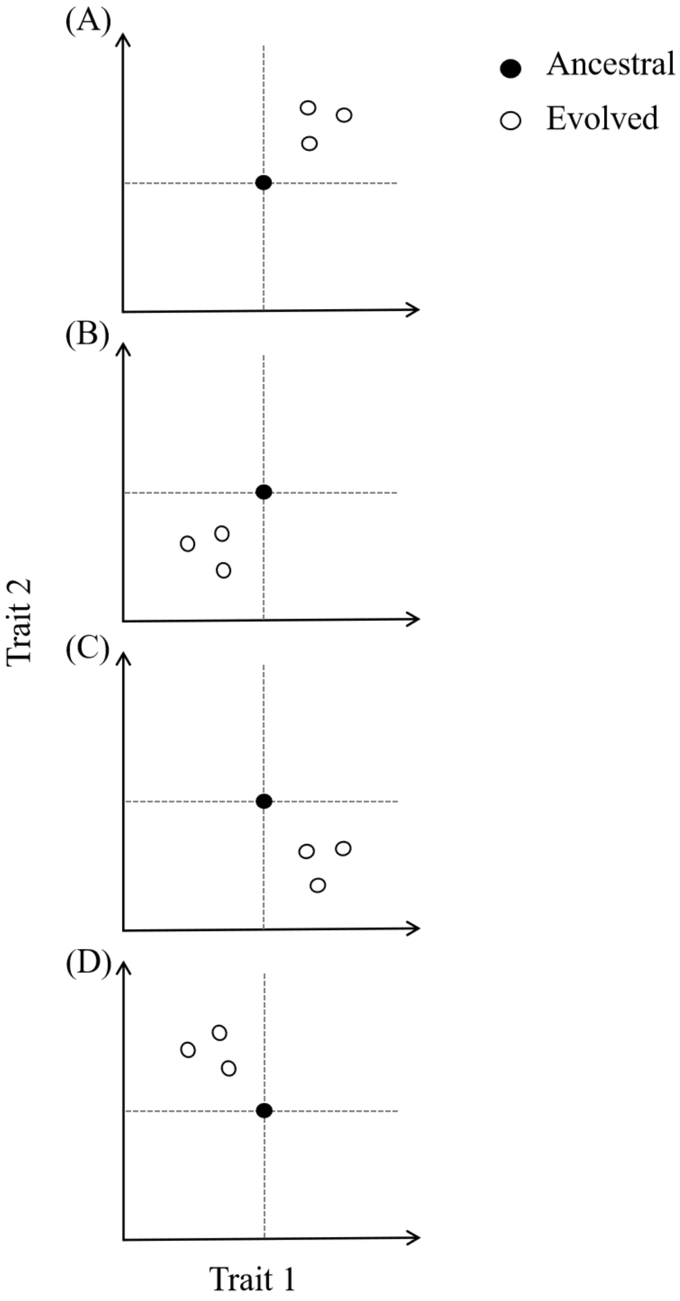

**Figure S2** The relationship between optical density ($OD_{750}$) and cell density of *Chlorella sorokiniana*. This relationship was established by making a gradient of algal culture dilutions, and measuring their cell densities using a microscope and $OD_{750}$ using a microplate reader.

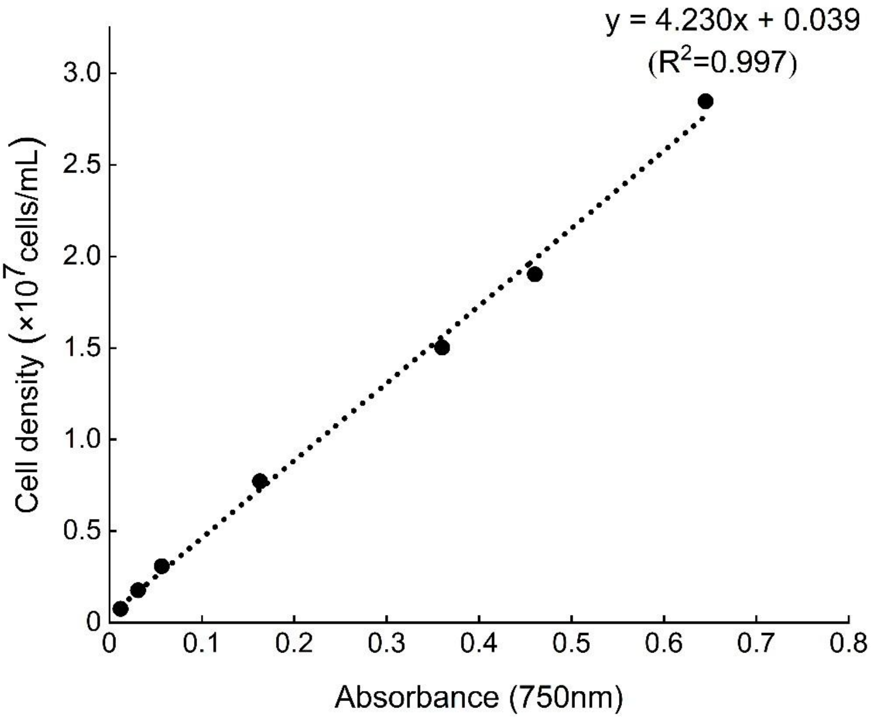

**Figure S3** Lipid standard curve was established based on the sulpho-phospho-vanillin (SPV) colorimetric method. Standard lipid stocks were prepared using commercially available soybean oil.

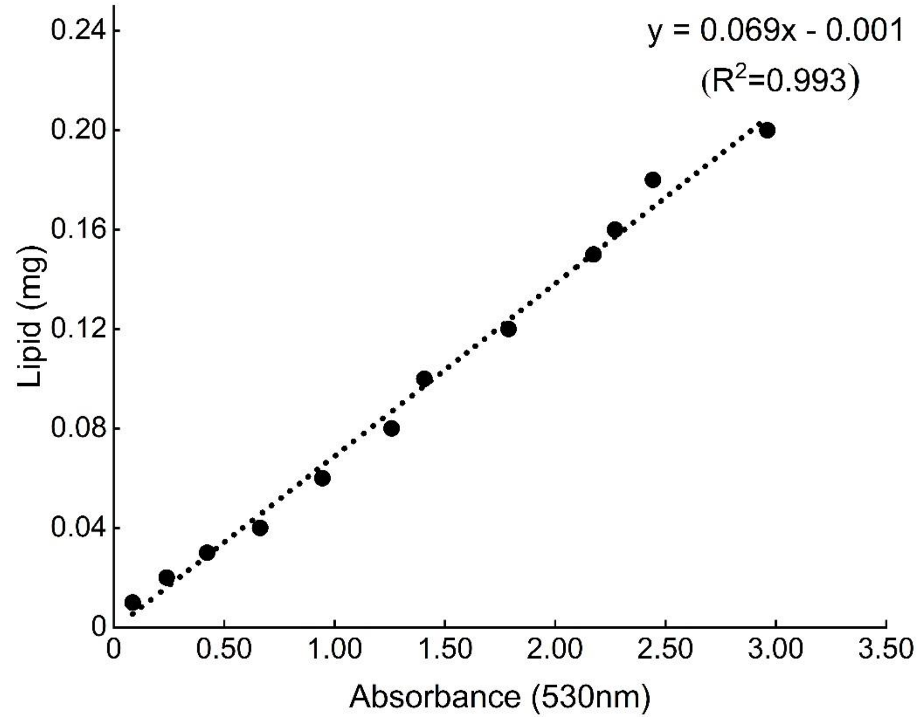

**Table S1** Summary of the selection response values in the 15°C selection lines. Here are shown the mean (± se) values of selection lines under each selection treatment (n = 4). Differences between the random-selection control and each of the other three treatments were analyzed using two-sample *t* tests.

| | Cell yield | | Lipid yield | | Lipid content per cell | |
|---|---|---|---|---|---|---|
| | Mean ± se | Difference from random-selection | Mean ± se | Difference from random-selection | Mean ± se | Difference from random-selection |
| Random | 0.59 ± 0.18 | | 0.06 ± 0.40 | | -0.53 ± 0.23 | |
| High-biomass | 0.46 ± 0.06 | *df* = 3.67, *P* = 0.535 | -0.13 ± 0.14 | *df* = 3.78, *P* = 0.673 | -0.59 ± 0.09 | *df* = 3.85, *P* = 0.806 |
| High-lipid | 0.54 ± 0.09 | *df* = 4.60, *P* = 0.815 | 0.12 ± 0.10 | *df* = 3.39, *P* = 0.887 | -0.41 ± 0.12 | *df* = 4.54, *P* = 0.689 |
| Rotation | 0.44 ± 0.10 | *df* = 4.86, *P* = 0.511 | 0.17 ± 0.26 | *df* = 5.12, *P* = 0.824 | -0.27 ± 0.15 | *df* = 5.21, *P* = 0.402 |

**Table S2** Summary of the selection response values in the 25°C selection lines. Here are shown the mean (± se) values of selection lines under each selection treatment (n = 4). Differences between the random-selection control and each of the other three treatments were analyzed using two-sample *t* tests.

| | Cell yield | | Lipid yield | | Lipid content per cell | |
|---|---|---|---|---|---|---|
| | Mean ± se | Difference from random-selection | Mean ± se | Difference from random-selection | Mean ± se | Difference from random-selection |
| Random | 0.60 ± 0.07 | | 1.21 ± 0.26 | | 0.61 ± 0.19 | |
| High-biomass | 0.62 ± 0.09 | *df* = 5.70, *P* = 0.874 | 1.62 ± 0.26 | *df* = 5.99, *P* = 0.315 | 0.99 ± 0.18 | *df* = 5.98, *P* = 0.202 |
| High-lipid | 0.67 ± 0.12 | *df* = 4.97, *P* = 0.636 | 1.50 ± 0.28 | *df* = 5.97, *P* = 0.478 | 0.83 ± 0.22 | *df* = 5.90, *P* = 0.479 |
| Rotation | 0.63 ± 0.08 | *df* = 5.86, *P* = 0.809 | 1.12 ± 0.42 | *df* = 5.05, *P* = 0.857 | 0.48 ± 0.35 | *df* = 4.68, *P* = 0.773 |

**Table S3** Summary of the measured values in the 15°C selection lines. Here are shown the mean (± se) values of selection lines under each selection treatment (n = 4). Differences between the ancestor and each selection treatment were analyzed using Welch's t tests or Wilcoxon test according to whether the data satisfies the assumptions of normal distribution and homogeneity of variance. Welch's t tests are performed because the response variable cell yield follows a normal distribution but does not meet the assumption of homogeneity of variance. Wilcoxon tests are performed because the response variables lipid yield and lipid content per cell do not meet the assumption conditions.

| | Cell yield ($\times 10^7$ cells $mL^{-1}$) | | Lipid yield (mg $L^{-1}$) | | Lipid content per cell (pg $cell^{-1}$) | |
|---|---|---|---|---|---|---|
| | Mean ± se | Difference from ancestor | Mean ± se | Difference from ancestor | Mean ± se | Difference from ancestor |
| Ancestor | 0.29 ± 0.01 | | 0.55 ± 0.19 | | 1.86 ± 0.65 | |
| Random | 0.55 ± 0.10 | $df = 3.02$, $P = 0.076$ | 0.73 ± 0.26 | W = 11, $P = 0.045$ | 1.20 ± 0.26 | W = 14, $P = 0.088$ |
| High-biomass | 0.47 ± 0.03 | $df = 3.19$, $P = 0.008$ | 0.50 ± 0.07 | W = 12, $P = 0.057$ | 1.05 ± 0.09 | W = 16, $P = 0.130$ |
| High-lipid | 0.51 ± 0.04 | $df = 3.08$, $P = 0.017$ | 0.63 ± 0.07 | W = 11, $P = 0.045$ | 1.27 ± 0.16 | W = 15, $P = 0.107$ |
| Rotation | 0.46 ± 0.04 | $df = 3.08$, $P = 0.030$ | 0.71 ± 0.14 | W = 10, $P = 0.035$ | 1.48 ± 0.19 | W = 13, $P = 0.071$ |
| Mean | 0.50 ± 0.03 | $df = 15.94$, $P < 0.001$ | 0.64 ± 0.07 | W = 44, $P = 0.002$ | 1.25 ± 0.09 | W = 58, $P = 0.008$ |

**Table S4** Summary of the measured values in the 25°C selection lines. Here are shown the mean (± se) values of selection lines under each selection treatment (n = 4). Differences between the ancestor and each selection treatment were analyzed using Welch's *t* tests or Wilcoxon test according to whether the data satisfies the assumptions of normal distribution and homogeneity of variance. Welch's *t* tests are performed because the response variable cell yield follows a normal distribution but does not meet the assumption of homogeneity of variance. Wilcoxon tests are performed because the response variables lipid yield and lipid content per cell do not meet the assumption conditions.

| | Cell yield (×$10^7$ cells $mL^{-1}$) | | Lipid yield (mg $L^{-1}$) | | Lipid content per cell (pg $cell^{-1}$) | |
|---|---|---|---|---|---|---|
| | Mean ± se | Difference from ancestor | Mean ± se | Difference from ancestor | Mean ± se | Difference from ancestor |
| Ancestor | 0.61 ± 0.02 | | 1.51 ± 0.39 | | 2.37 ± 0.55 | |
| Random | 1.13 ± 0.08 | $df = 3.28$, $P = 0.007$ | 5.64 ± 1.56 | W = 1, $P = 0.003$ | 4.79 ± 0.97 | W = 4, $P = 0.007$ |
| High-biomass | 1.16 ± 0.10 | $df = 3.18$, $P = 0.012$ | 8.40 ± 2.26 | W = 0, $P = 0.002$ | 7.02 ± 1.46 | W = 2, $P = 0.004$ |
| High-lipid | 1.23 ± 0.16 | $df = 3.07$, $P = 0.030$ | 7.60 ± 1.98 | W = 1, $P = 0.003$ | 6.02 ± 1.05 | W = 3, $P = 0.005$ |
| Rotation | 1.16 ± 0.10 | $df = 3.21$, $P = 0.009$ | 5.85 ± 2.06 | W = 4, $P = 0.007$ | 4.72 ± 1.40 | W = 7, $P = 0.017$ |
| Mean | 1.17 ± 0.05 | $df = 18.43$, $P < 0.001$ | 6.87 ± 0.94 | W = 6, $P < 0.001$ | 5.64 ± 0.61 | W = 16, $P < 0.001$ |
